# Collagen and actin network mediate antiviral immunity against Orsay in C. elegans intestinal cells

**DOI:** 10.1101/2023.04.20.537671

**Authors:** Ying Zhou, Weiwei Zhong, Yizhi Jane Tao

## Abstract

C. elegans is a free-living nematode that is widely used as a small animal model for studying fundamental biological processes and disease mechanisms. Since the discovery of the Orsay virus in 2011, C. elegans also holds the promise of dissecting virus-host interaction networks and innate antiviral immunity pathways in an intact animal. Orsay primarily targets the worm intestine, causing enlarged intestinal lumen as well as visible changes to infected cells such as liquefaction of cytoplasm and rearrangement of the terminal web. Previous studies of Orsay identified that C. elegans is able to mount antiviral responses by DRH-1/RIG-I mediated RNA interference and Intracellular Pathogen Response, a uridylyltransferase that destabilizes viral RNAs by 3′ end uridylation, and ubiquitin protein modifications and turnover. To comprehensively search for novel antiviral pathways in *C. elegans*, we performed genome-wide RNAi screens by bacterial feeding using existing bacterial RNAi libraries covering 94% of the entire genome. Out of the 106 antiviral genes identified, we investigated those in three new pathways: collagens, actin remodelers, and epigenetic regulators. By characterizing Orsay infection in RNAi and mutant worms, our results indicate that collagens likely form a physical barrier in intestine cells to inhibit viral infection by preventing Orsay entry. Furthermore, evidence suggests that the intestinal actin (*act-5*), which is regulated by actin remodeling proteins (*unc-34*, *wve-1* and *wsp-1*), a Rho GTPase (*cdc-42*) and chromatin remodelers (*nurf-1* and *isw-1*), also provides antiviral immunity against Orsay possibly through another physical barrier presented as the terminal web.

## INTRODUCTION

Since its first discovery in 2011, Orsay remains the only virus known to naturally infect the small animal model *Caenorhabditis elegans* (*C. elegans*) (1). Based on phylogenetic analysis, Orsay and three other subsequently discovered viruses Le Blanc, Santeuil and Melnik, all of which infect *Caenorhabditis briggsae* (*C. briggsae*), form a close cluster that is distantly related to members of the *Nodaviridae* family (2). All four nematode viruses transmit through the oral-fecal route with tissue tropism towards the worm intestine. Infected animals exhibit enlarged intestinal lumen that is accompanied by subcellular structural changes such as reorganization of the terminal web, diminished intermediate filament, and liquefaction of the cytoplasm (1). Orsay infection results in reduced food consumption and a smaller body size, but does not affect animal life span or brood size.

Orsay has a positive-sense, bi-segmented RNA genome of ∼6.3 kb that encodes three open reading frames (ORFs), including the putative RNA-dependent RNA polymerase (RdRP), the viral capsid protein (CP), and a nonstructural protein δ (1). In addition, a CP-δ fusion protein is produced during virus infection through ribosomal frameshift, which is mediated by a RNA stem loop conserved in nematode viruses (3). Orsay capsid has a T=3 icosahedral symmetry with trimeric protrusions formed by the C-terminal region of the CP polypeptide (4). Despite the lack of detectable sequence identity, the structural fold of Orsay CP resembles that of the betanodavirus virus capsid protein (5). It has been shown that the free δ and the CP-δ fusion protein play different roles during viral exit and viral entry, respectively. While free δ was found to colocalize with terminal web and mediate nonlytic viral release (6), CP-δ is incorporated into infectious viral particles as a pentameric head fiber (7). The CP-δ head fiber adopts a novel β-bracelet structure with a terminal globular domain that is likely involved in host receptor engagement. A system for engineering recombinant Orsay virus is available utilizing transgenic worms expressing the two viral RNA segments (8).

Considering the simplicity of the Orsay virus and the extensive genetic and molecular tools available for *C. elegans* as a laboratory model, the pair presents a promising opportunity for studying virus-host interactions and innate immune responses. Due to the lack of adaptive immunity, *C. elegans* uses exclusively innate immune responses to defend against viral and microbial infections. The intestine of *C. elegans* is comprised of 20 large epithelial cells that are positioned as bilaterally pairs forming a long tube with a central lumen. These intestinal cells exhibits molecular polarity similar to mammalian intestines in terms of extracellular matrix and apical membrane surface with underlying terminal web (9). Importantly, genomic comparison of humans and *C. elegans* demonstrated that the majority of human disease genes and disease pathways are present in the worm (10). Among the available 18,452 *C. elegans* protein sequences, at least 83% has homologous genes in human (11). Many genes involved in defense against bacterial pathogens in *C. elegans* also function in humans as host defense genes (12).

Studies of Orsay infection in the past few years have uncovered several conserved antiviral defense mechanisms. These include the RNA interference pathway that degrades viral RNA and triggers antiviral gene expression (1, 13, 14); an inductive transcriptional response called Intracelluar Pathogen Response (15); a uridylyltransferase that destabilizes viral RNAs by 3′ end uridylation (16); and ubiquitin protein modifications and turnover (17, 18). In another study, bioinformatics analyses of expressed genes during Orsay infection revealed the presence of uncharacterized anti-stress pathways (19). In addition to antiviral genes, a forward genetic screen implicated several genes and their human orthologs in endocytosis (i.e. SID-3 and WASP) to be required for viral infection (20). In spite of the progress made, the fact that only a few above-mentioned pathways have been linked to antiviral immunity shows that our understanding of the antiviral responses of *C. elegans* is still limited.

To comprehensively search for novel antiviral pathways in *C. elegans*, we performed genome-wide RNAi screens by bacterial feeding using existing bacterial RNAi libraries covering 94% of the entire genome. We identified at least 106 antiviral genes, including those belonging to known antiviral mechanisms as well as those having not been associated with *C. elegans* antiviral immunity. We further investigated those in three new pathways: collagens, actin remodelers, and epigenetic regulators. By characterizing Orsay infection in RNAi and mutant worms, our results indicate that collagens, including *col-51*, *col-61*, *col-92*, *cutl-21*, and *sqt-2*, likely form a physical barrier in intestine cells to inhibit viral infection by preventing Orsay entry. Additionally, evidence suggests that the intestinal actin (*act-5*), which is regulated by actin remodeling proteins (*unc-34*, *wve-1* and *wsp-1*), a Rho GTPase (*cdc-42*) and chromatin remodelers (*nurf-1* and *isw-1*), also provides antiviral immunity against Orsay possibly through another physical barrier presented as the terminal web.

## RESULTS

### A genome-wide RNAi screen for antiviral genes

We designed a genome-wide RNAi screen to discover which genes are required for *C. elegans* antiviral immunity. We started with a strain that is resistant to the Orsay virus. Starting from the first larval stage (L1), these worms were fed with RNAi bacteria to inactivate various worm genes, and infected with Orsay. We examined the severity of infection when the worms were day-3 adults (Fig. 1A). The worms also had the temperature-sensitive mutation *glp-4(bn2ts)* so that they were sterile at 20°C, allowing us to grow them to day-3 adults without the interference from any progeny. The rationale of the screen was that without RNAi, worms would be resistant to Orsay infection and asymptomatic; RNAi inactivation of an antiviral gene would make the worms sensitive to viral infection and display the infection symptom of transparent intestine (Fig. 1A). To ensure that the transparent intestine phenotype was specific to viral infection and not caused by RNAi, we also examined RNAi worms without viral infection to ensure that these worms did not have the transparency phenotype (Fig. 1A).

**Figure 1.**
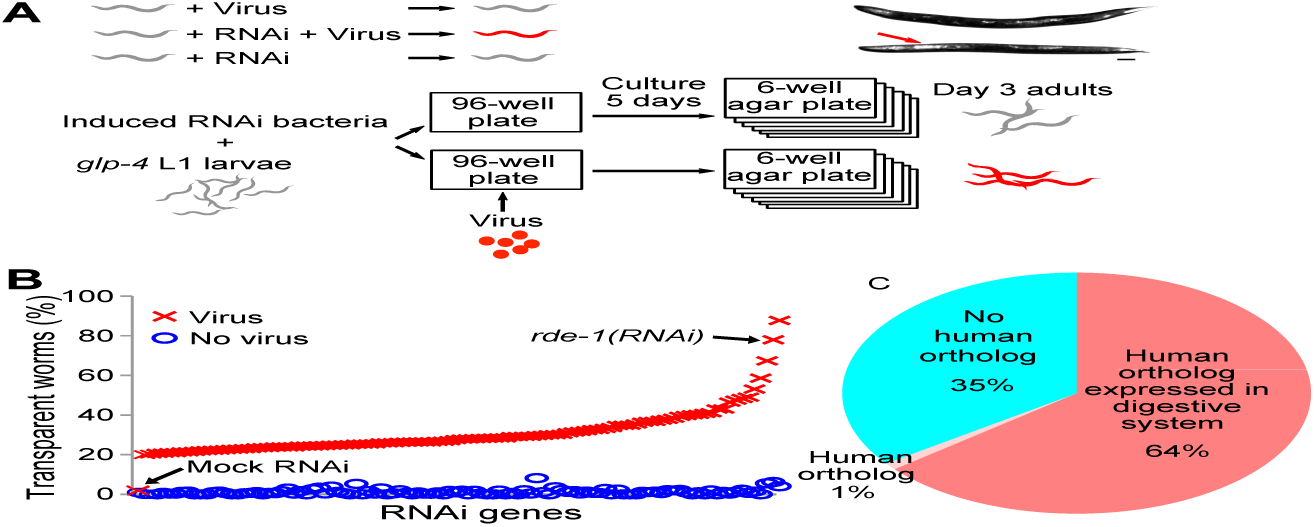
Genome-wide RNAi screen for antiviral genes. A) Schematic drawing of the design of the antiviral gene screen. A strain resistant to Orsay infection was used. We screened for RNAi that made the strain sensitive to Orsay infection. Top right image, the transparent anterior intestine phenotype was scored as infection symptom. Scale bar, 100μm. Uninfected RNAi worms were examined to ensure that the transparent intestine phenotype was caused by virus infection and not just RNAi. B) Percentage of worms with transparent intestines after RNAi of antivirus genes. *rde-1* and mock RNAi were positive and negative controls, respectively. C) Fractions of worm antivirus genes with human orthologous genes.

Bacteria containing the empty vector were used as the negative control. As expected, very few worms (< 2%) in the negative control group showed the transparency phenotype with or without the virus (Fig. 1B). *rde-1(RNAi)* bacteria were used as the positive control. In this group, 77.9% of infected *rde-1(RNAi)* worms showed the transparency phenotype while only 2.8% of uninfected *rde-1(RNAi)* worms did, showing a difference of 75.1% (Fig. 1B). The difference between the percentage of transparent worms in the virus-infected group and the uninfected group was used to select hit genes, and a threshold of 20% was applied. Including the positive control *rde-1*, a total of 106 genes were identified as hits (Fig. 1B, Table S1). RNAi of these genes reproducibly displayed elevated levels of transparent worms upon viral infection, suggesting that these genes are required for antiviral immunity.

Most of these antiviral genes have human orthologs. 69 out of these 106 antiviral genes (65%) have orthologous genes in human (WormBase W277, Fig. 1C). Since Orsay infects intestine cells, we examined how many of these orthologous genes were expressed in the human digestive system. Almost all (68/69) genes with human orthologs have orthologs expressed in the human digestive system (Fig. 1C).

These antiviral genes cover a broad range of functions (Table S1). We discovered several genes belonging to known antiviral mechanisms such as RNAi pathways (*e.g.*, *rsd-6*) and ubiquitin-mediated protein degradation machinery (*e.g.*, *fbxb-85*, *nhl-1*, *rnf-113, usp-48*). More importantly, we discovered genes in pathways that have not been associated with *C. elegans* antiviral mechanisms, such as transcription regulation genes (*e.g.*, *attf-6, npr-34, taf-9, thoc-2, unk-1, ztf-28*), and RNA processing genes (*e.g., edc-4, prpf-4*). We further investigated three of these new pathways: collagens, actin remodelers, and epigenetic regulators, and report our findings below.

### Collagens mediate antiviral innate immunity

RNAi of five collagens, *col-51*, *col-61*, *col-92*, *cutl-21*, and *sqt-2*, significantly increased the number of worms with the symptom of transparent intestine upon Orsay infection (Fig. 2A).

**Figure 2.**
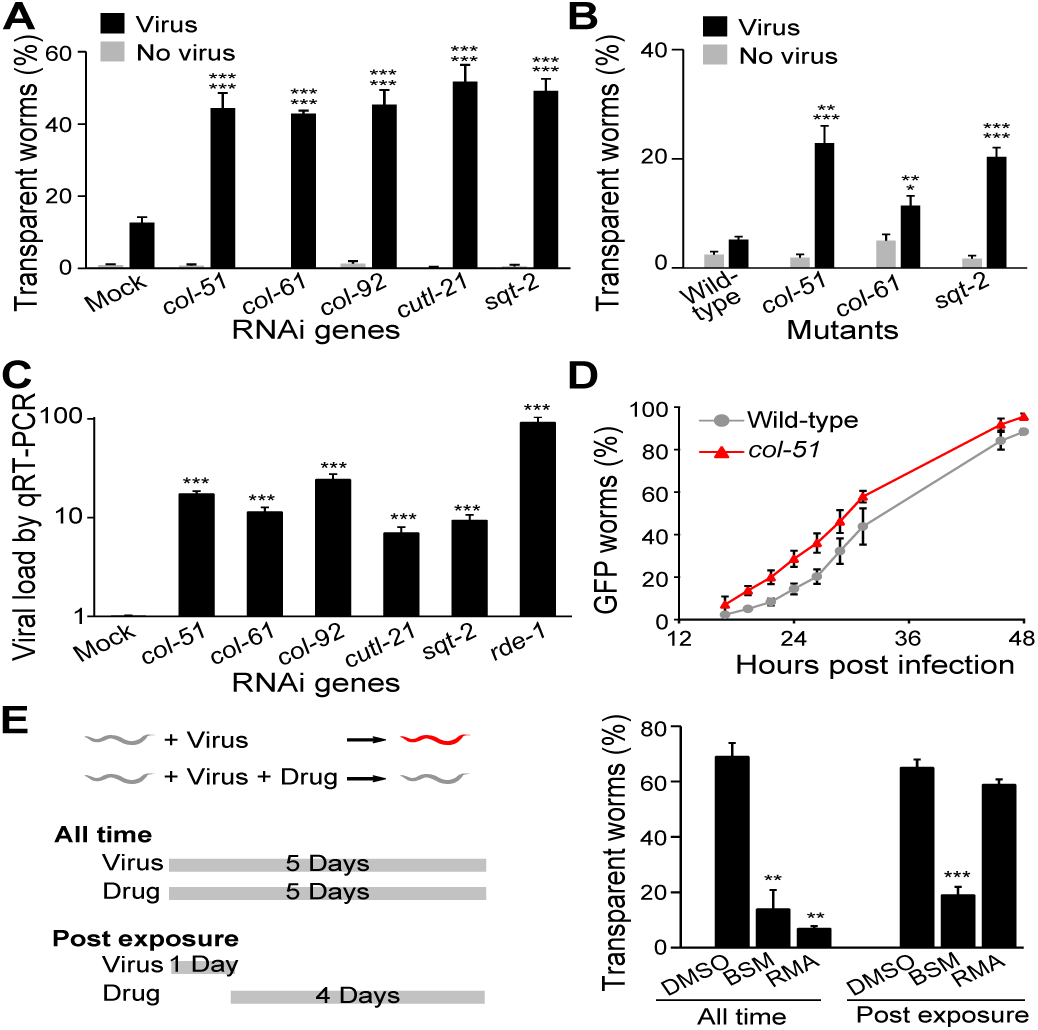
Collagens are required for antiviral immunity. A, B) Percentage of worms with transparent intestines after collagen RNAi (A) or in a collagen mutant (B). P value comparing virus vs. no virus groups of the same genotype was labeled as the first line, while p value comparing virus groups of a given genotype vs. mock or wild-type was labeled as the second line. n ≥ 6 plates for all groups. C) Viral load detected by qRT-PCR in infected worms upon various collagen RNAi. At least three independent experiments were tested with at least three technical replicates in each experiment. D) Infection dynamics as measured by *Ppals-5::GFP* worms showing reporter GFP. n ≥ 3 plates for all data points. E) Antiviral drug effects. A strain sensitive to Orsay infection was used. Worms were dosed with resorcinol monoacetate (RMA), bismuth subsalicylate (BSM), or the solvent DMSO, and exposed to virus using two exposure methods. Left, schematic drawing of the two exposure methods. Right, Percentage of transparent worms in all test groups. Bars and error bars indicate means and standard errors. *, p < 0.05, ** p < 0.01, ***, p<0.001, Student’s t-test.

Among these five genes, three (*col-51*, *col-61*, and *sqt-2*) have homozygous-viable mutants publicly available. These mutants displayed the same phenotype of increased number of symptomatic worms upon viral infection (Fig. 2B), confirming that these genes mediate antiviral immunity. In addition, we applied qRT-PCR to examine the viral load in the RNAi worms. In this experiment, N2 worms without the *glp-4* mutation were used to eliminate any doubts on *glp-4* effects. Consistent with the phenotype of increased symptomatic worms, RNAi of these five collagens significantly increased viral load in worms (Fig. 2C). Altogether, these data demonstrated that these collagens are required for antiviral immunity.

### Collagen-mediated antiviral immunity functions at an early stage of viral infection

An intuitive hypothesis for the collagen antiviral function is that collagens may act as an exterior barrier blocking the Orsay virus from entering host cells. To test that, the reporter strain *Ppals-5::GFP* was used to monitor whether collagen inactivation changed viral infection dynamics. This reporter turns on GFP expression upon Orsay virus infection (21). We constructed strains with the reporter on either the wild-type N2 or the *col-51* mutant background, cultured these worms, added the Orsay virus when they reached the L4 larval stage, and examined every two hours for the reporter GFP expression. In comparison with the wild-type N2 worms, more *col-51* worms showed GFP at the same time point (Fig. 2D). Overall, the *col-51* mutation shifted the infection dynamics curve to about four hours earlier (Fig. 2D), consistent with the hypothesis of collagens as a viral entry barrier.

Since inactivation of collagens reduced antiviral immunity, we asked whether enhancing collagens would increase antiviral immunity. To test that, we used the *drh-1;glp-4* mutant strain, which is highly sensitive to Orsay infection. We tested the effect of resorcinol monoacetate (RMA), a chemical that crosslinks collagens (22), on these worms. Indeed, when worms were exposed to Orsay and RMA simultaneously, significantly fewer worms showed the infection symptom of transparent intestine than the group without RMA (Fig. 2E).

Interestingly, RMA was only effective when applied at an early stage of viral infection. In a post exposure application experiment where the worms were exposed to the virus first for one day and then exposed to the drug for four days, RMA showed no protective effect (Fig. 2E). As a control, the commonly used antidiarrheal drug bismuth subsalicylate (BSM) displayed the same protective effects in both early application and post exposure application experiments (Fig. 2E). The early requirement for RMA effects suggested that collagen-mediated antiviral immunity functions at an early stage of viral infection, such as viral entry.

### Collagens function in intestine cells to mediate antiviral immunity

Since Orsay only infects intestine cells (23), we questioned whether the site-of-action for the antiviral collagens was intestine cells. We applied collagen RNAi on a tissue-specific RNAi strain that was only sensitive to RNAi in intestine cells. This strain, MW265, had the RNAi-resistant mutation of *rde-1*. The intestine cells were restored to be RNAi sensitive by the rescue transgene of *rde-1* driven by the intestine-specific *ges-1* promoter. This strain was resistant to Orsay virus infection (Fig. 3A). Inactivation of the collagens in intestine cells by RNAi significantly increased the virus sensitivity of this strain (Fig. 3A), suggesting that collagen-mediated antiviral immunity was needed in intestine cells.

**Figure 3.**
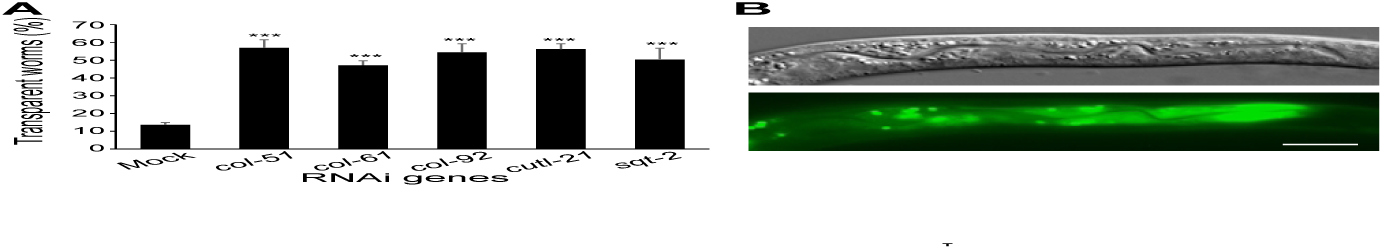
Antiviral collagens function in the intestine. A) Percentage of transparent worms in Orsay-infected worms after intestine-specific RNAi of various collagens. Bars and error bars indicate means and standard errors. ***, p<0.001 comparing to mock RNAi, Student’s t-test. B) *col-51* GFP expression in intestine cells of a *Pcol-51::col-51::GFP* worm. Scale bar, 5 μm.

*C. elegans* has two types of collagens: cuticle collagens and type IV basement membrane collagens (24). No collagens have been reported to be synthesized in intestine cells: Cuticle collagens are believed to be synthesized in hypodermis (25), whereas the type IV collagens are synthesized in muscles and somatic gonads (26). Therefore, it was a surprising discovery that the site-of-action for these antiviral collagens was intestine cells. To confirm that these antiviral collagens were expressed in intestine cells, we constructed a translational fusion of *Pcol-51::col-51::GFP* and made a transgenic worm strain with this reporter. The reporter showed that *col-51* was expressed in intestine cells in all developmental stages from L1 to adults, with the highest expression observed during early larval stages. During these early larval stages, *col-51* expression was observed in all intestine cells, with the highest expression in posterior intestine cells (Fig. 3B). Altogether, these data demonstrated that there are collagens generated by intestine cells and that these intestinal collagens mediate antiviral innate immunity.

### Actin remodelers mediate antiviral innate immunity

Several genes encoding actin-remodeling proteins were also identified as antiviral genes in our RNAi screen. RNAi inactivation of *wsp-1* and *wve-1*, two genes encoding the evolutionarily conserved actin regulators WSP-1/WASP and WVE-1/WAVE, respectively, significantly increased the percentage of worms showing the viral infection symptom of transparent intestine (Fig. 4A). A heterozygous *wve-1* mutant confirmed the RNAi phenotype (Fig. 4B). However, a homozygous *wsp-1* mutant (*gm324)* did not show such enhanced infection phenotype (Fig. 4B). The discrepancy between this mutant and RNAi is likely due to the fact that the *wsp-1* mutant allele *gm324* only affects one of the two *wsp-1* isoforms (Fig. 4C). Our *wsp-1* RNAi was efficient as it reduced nearly 80% of the total *wsp-1* transcript level (Fig. 4D). RNA inactivation of *cdc-42*, a gene encoding the ortholog of the mammalian WASP regulator CDC42, showed similar effects of increased infection symptoms (Fig. 4A), and such phenotype was observed in a heterozygous *cdc-42* mutant (Fig. 4B). Our data suggested that these proteins that are known to regulate actin dynamics are crucial for antiviral immunity in *C. elegans*.

**Figure 4.**
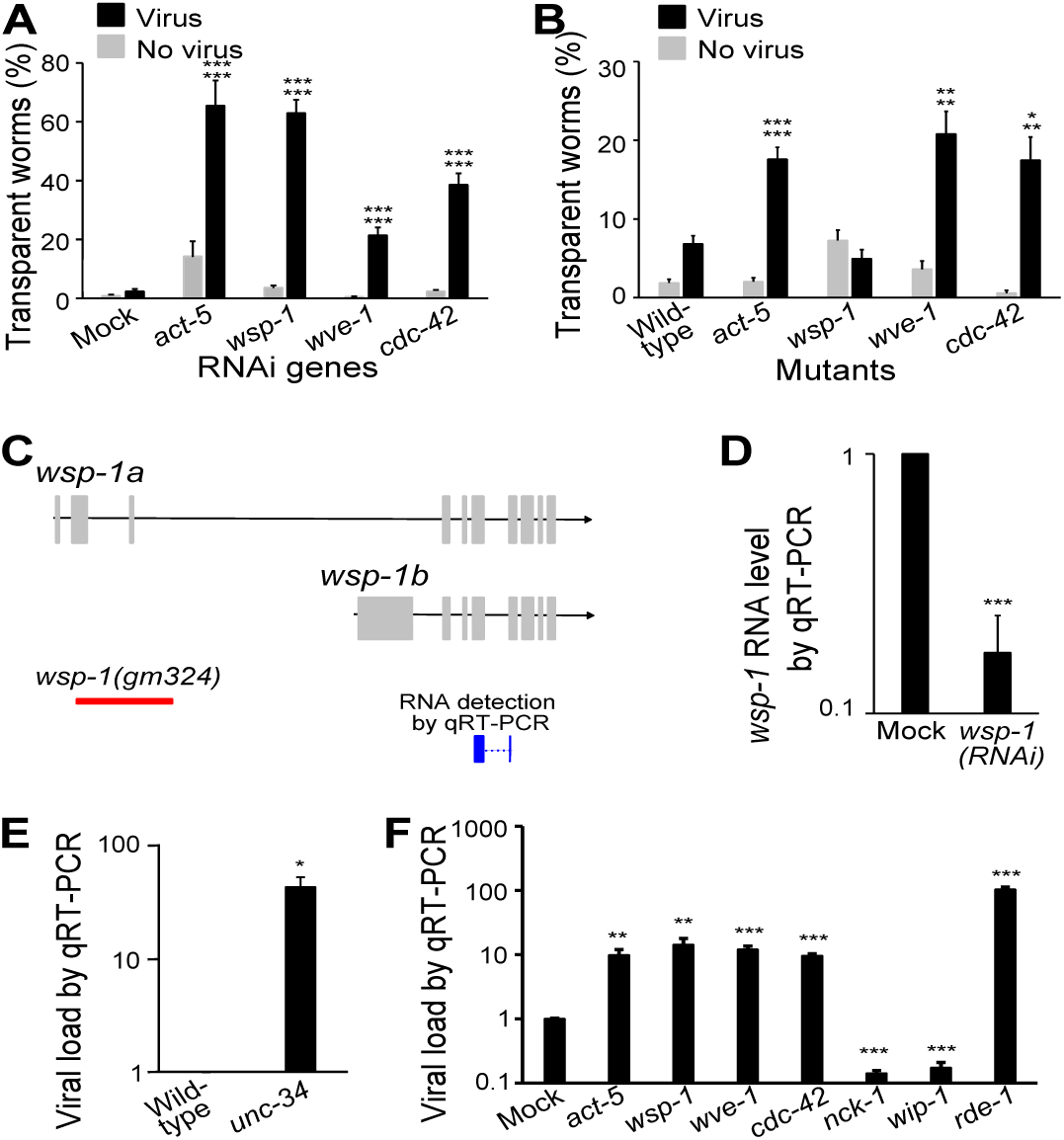
Actin remodelers are required for antiviral immunity. A, B) Percentage of worms with transparent intestines after RNAi (A) or in a mutant (B). P value comparing virus vs. no virus groups of the same genotype was labeled as the first line, while p value comparing virus groups of a given genotype vs. mock or wild-type was labeled as the second line. n ≥ 6 plates for all groups. C) Schematic drawing of *wsp-1* transcripts, area of deletion in the allele *gm324*, and region of detection in our qRT-PCR. D) *wsp-1* RNA level in *wsp-1(RNAi)* worms as determined by qRT-PCR. E) Viral load determined by qRT-PCR in infected *unc-34* mutants. F) Viral load detected by qRT-PCR in infected worms upon RNAi of actin and various actin modulators. Bars and error bars indicate means and standard errors. *, p < 0.05, ** p < 0.01, ***, p<0.001, Student’s t-test.

In addition to *WSP-1* and *WVE-1*, *UNC-34*/Ena/VASP is another known actin remodeler (27). Since *unc-34* was not in our RNAi library, we examined a homozygous *unc-34* mutant. The mutant showed a transparent intestine phenotype even without viral infection, so we used qRT-PCR to examine the viral load and found that the viral load was significantly increased in this mutant (Fig. 4E). Therefore, similar to *wsp-1* and *wve-1*, *unc-34* also mediates antiviral immunity in *C. elegans*.

These results led us to hypothesize that actin is involved the *C. elegans* antiviral defense. In *C. elegans*, the actin isoform ACT-5 is intestine specific and is an essential component of the terminal web (26). Since *act-5* RNAi causes larval lethality, we diluted *act-5* RNAi with control bacteria at a ratio of 1:50 to reduce its effectiveness so that the RNAi worms could grow to adults. With this reduced RNAi, we were able to test our hypothesis. Indeed, *act-5(RNAi)* significantly increased the percentage of worms displaying the infection symptom (Fig. 4A), confirming that actin mediates antiviral immunity in the intestine cells. Similar phenotype was confirmed in a heterozygous *act-5* mutant (Fig. 4B).

The increased viral infection phenotype upon RNAi of *act-5*, *wsp-1*, *wve-1*, and *cdc-42* was also confirmed by measuring viral load with qRT-PCR. As expected, RNAi of these genes caused a significant increase in viral load (Fig. 4F). Interestingly, RNAi of two other WASP regulators, *nck-1* and *wip-1*, did not increase the viral load. Instead, they significantly reduced the viral load (Fig. 4F), suggesting that different actin remodelers have different functions during the course of viral infection.

### Epigenetic regulation is involved in mediating antiviral innate immunity

Several genes encoding proteins regulating gene expression showed as hits in our antiviral genetic screen, for example, genes encoding the histone acetyltransferase *MYS-1*, the nucleosome remodeling factor *NURF-1*, and the chromatin regulator *ISW-1*. We examined the mutants of two of these genes, *nurf-1* and *isw-1*. RNAi of these genes significantly increased the percentage of symptomatic animals upon infection (Fig. 5A). Similarly, homozygous mutants of these genes also showed an increased percentage of symptomatic animals (Fig. 5B). Consistently, when we examined the viral load by qRT-PCR upon infection, these mutants had significantly higher viral load than the wild-type N2 worms (Fig. 5C). The requirement of chromatin remodelers *NURF-1* and *ISW-1* in antiviral defense showed that epigenetic regulation is a crucial part of the host antiviral innate immune response.

**Figure 5.**
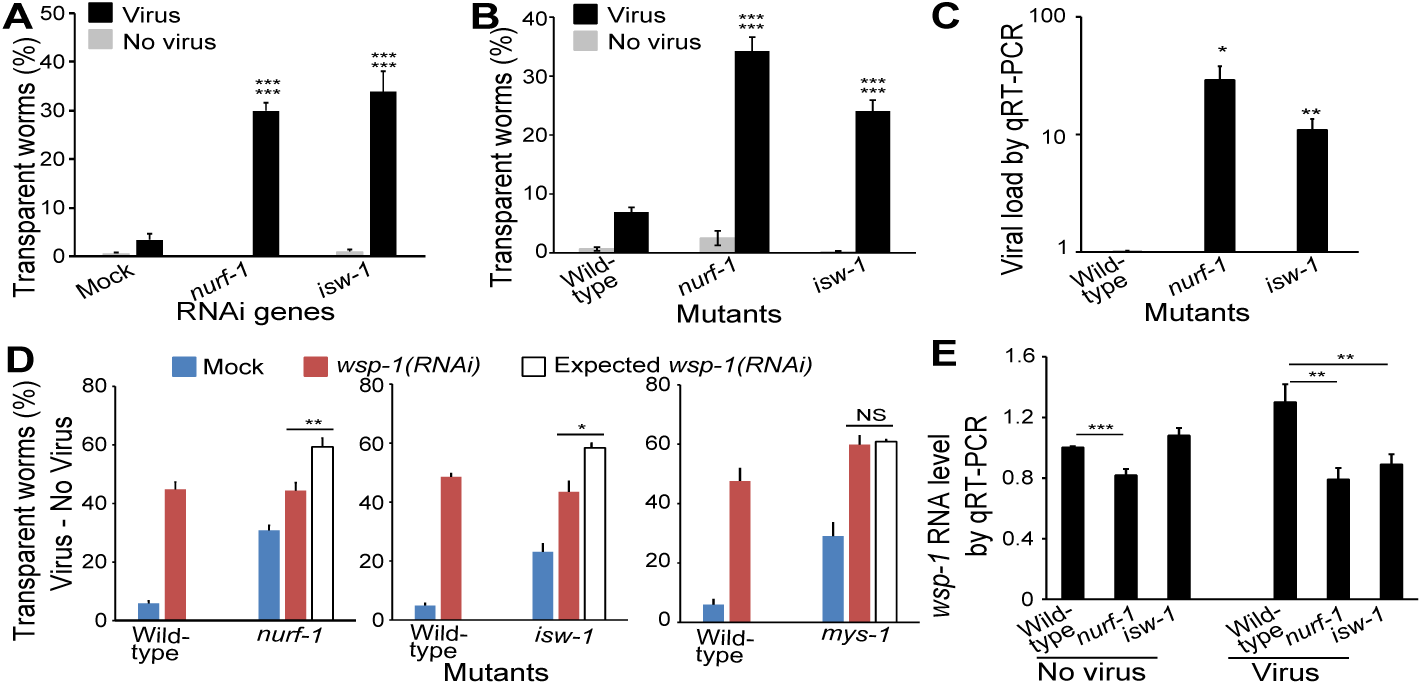
Chromatin remodelers are required for antiviral immunity. A, B) Percentage of worms with transparent intestines after RNAi (A) or in a mutant (B). P value comparing virus vs. no virus groups of the same genotype was labeled as the first line, while p value comparing virus groups of a given genotype vs. mock or wild-type was labeled as the second line. C) Viral load detected by qRT-PCR in infected mutants. D) Percentage of worms with infection caused symptoms in various genotypes. Unfilled bar indicated expected rate of symptom if mutation and RNAi genes had additive effects (i.e., no epistatic interaction). E) *wsp-1* RNA level detected by qRT-PCR. Bars and error bars indicate means and standard errors. NS, not significant, *, p < 0.05, ** p < 0.01, ***, p<0.001, Student’s t-test.

### Chromatin remodelers *NURF-1* and *ISW-1* function through the actin remodeler *WSP-1* to mediate antiviral innate immunity

*NURF-1* and *ISW-1* was reported to regulate the expression of *wsp-1* to modulate axon guidance during *C. elegans* development (28). Since all three genes showed as antiviral genes in our screen, we tested whether *wsp-1* is also the target of *nurf-1* and *isw-1* in antiviral defense using quantitative epistasis analysis. We performed *wsp-1(RNAi)* on both wild-type N2 worms and *nurf-1* mutant worms, scored the percentage of symptomatic worms, and applied a quantitative epistasis model to calculate the expected phenotypic severity of two-gene inactivation based on additive effects of single gene inactivation (Fig. 5D). *wsp-1(RNAi);nurf-1* worms had significantly less severe phenotype than the expected value, suggesting a genetic interaction between *wsp-1* and *nurf-1* in antiviral functions (Fig. 5D). If *nurf-1* and *wsp-1* function independently, then the percentage of symptomatic worms should be higher in the population with two genes inactivated than the population with a single gene inactivated. In contrast, we found that *wsp-1(RNAi)* on *nurf-1* mutants and wild-type worms had the same percentage of symptomatic worms (Fig. 5D, 44.3±2.8% vs. 44.8±2.5%, p = 0.9) despite that *nurf-1* mutants had more symptomatic worms than wild-type worms (Fig. 5D, 30.8±1.7% vs. 5.8±0.9%, p < 0.0001). These data supported the hypothesis that *nurf-1* regulates *wsp-1* expression to mediate antiviral response. Analysis on *wsp-1(RNAi)* on *isw-1* mutants showed similar results (Fig. 5D). As a control, analysis on *wsp-1* and another chromatin remodeler *mys-1* showed that *wsp-1(RNAi);mys-1* worms had the same percentage of symptomatic worms as the expected value (Fig. 5D, 59.9±2.9% vs. 50.8±0.7%, p = 0.7), demonstrating that *mys-1* does not act through *wsp-1*. Together, these results suggest that *nurf-1* and *isw-*1, but not *mys-1*, regulate *wsp-1* expression to modulate antiviral immunity.

To directly test the impact of *nurf-1* and *isw-*1 mutation on *wsp-1* RNA levels, we quantified *wsp-1* RNA levels in these mutants by qRT-PCR under both infected and uninfected conditions. In the uninfected group, the *wsp-1* RNA level difference between mutants and wild-type animals were less notable, with *wsp-1* RNA being lower only in *nurf-1* mutants (Fig. 5E). Upon viral infection, wild-type animals significantly increased its *wsp-1* RNA level from the uninfected level (Fig. 5E, 1±0.01 vs. 1.3±0.12, p < 0.05), but the levels of *wsp-1* RNA in mutant worms remained essentially unchanged. As a result, in the virus-infected group, *wsp-1* RNA levels in both *nurf-1* and *isw-*1 mutants were significantly lower than that in wild-type animals (Fig. 5E). These data suggested that *nurf-1* and *isw-*1 are required to up-regulate *wsp-1* expression upon viral infection. While the *wsp-1* RNA level changes were significant, the magnitude of changes was rather mild (Fig. 5E). This was consistent with previous report on *wsp-1* RNA level changes in these mutants during *C. elegans* development (28). Several factors could contribute to this low magnitude of changes. For example, the *wsp-1* RNA changes may be limited to intestine cells, therefore quantifying whole-body *wsp-1* RNA levels would record a reduced magnitude of changes.

## DISCUSSION

Our genome-wide *C. elegans* RNAi screen revealed 106 antiviral genes of diverse functions. The diversity of gene functions among these antiviral genes demonstrated that *C. elegans* has complex innate antiviral mechanisms. By implementing multiple controls and applying a stringent cutoff, our RNAi screen prioritized reducing false positives over false negatives. As a result, the screen is not saturated. For example, we did not recover known antiviral genes such as *pals-22* (29) and *cde-1* (*16, 30*). More studies are needed to identify a comprehensive list of antiviral genes.

Our investigation of antiviral genes encoding collagens, actin remodelers, and epigenetic regulators suggested the following antiviral model of building physical barriers for the Orsay virus (Fig. 6). The first barrier includes collagens generated by the intestine cells (Fig. 6). Similar to the human intestine epithelium, *C. elegans* intestine also has a glycocalyx layer on the apical side of microvilli within the digestive tract (26). These collagens and possibly other glycoproteins can function in the glycocalyx to form a physical barrier to block the viral entry to intestine cells. In support of this model, our data showed that intestine cells make collagens (Fig. 3). Treating worms with a collagen crosslinking drug RMA delayed the Orsay infection time course by at least four hours (Fig. 2), suggesting that these collagens mediate antiviral functions at an early stage of infection such as viral entry.

**Figure 6.**
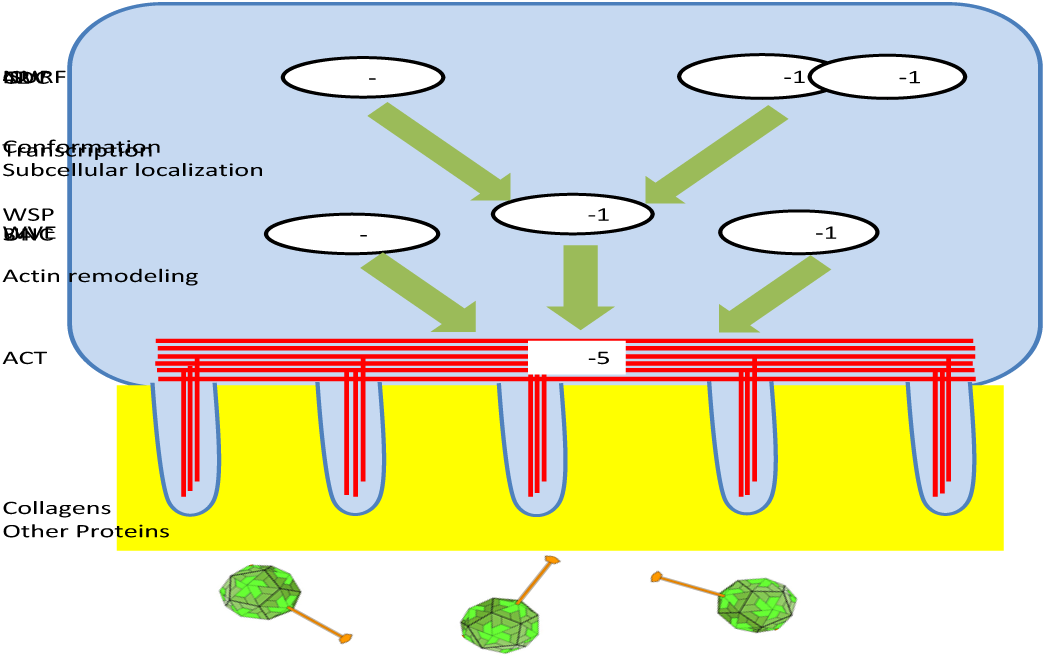
Model of *C. elegans* physical barrier based innate immunity against the Orsay virus. Collagens and other proteins in the glycocalyx is the first layer of physical barrier against the virus. ACT-5 actin in the terminal web is the second layer to block the virus. The intestine cells regulate actin through actin remodeling proteins UNC-34, WSP-1, and WVE-1. WSP-1 activity is modulated by CDC-42 while its expression is regulated by NURF-1 and ISW-1 to coordinate the antiviral defense.

The intestinal terminal web composed mostly of actin and intermediate filaments likely constitutes another physical barrier for the Orsay virus. Based on our results, we propose that actin remodelers *UNC-34*, *WSP-1*, *WVE-1* reorganize ACT-5 in terminal web to mediate such antiviral defense (Fig. 6). It was previously shown that CDC42, a Rho GTPase, regulates the membrane localization of WSP-1 which then promotes the polymerization of F-actin (31). Likewise, here we demonstrated that *WSP-1* activity is regulated by *CDC-42* in *C. elegans*. Additionally, *WSP-1* transcription is upregulated by chromatin remodelers *NURF-1* and *ISW-1* (Fig. 6). All these actin-regulating factors are required for antiviral immunity (Figs. 4, 5), suggesting that this is a highly coordinated activity. Although our data strongly implicate actin/actin modelers in antiviral immunity, we cannot yet determine whether the antiviral effect occurs during viral entry, viral release, or at other stages of the infection life cycle. Because our transparency assay and RT-qPCR analysis for scoring Orsay infection were performed at five and two days after infection, respectively, multiple rounds of replication could have taken place and therefore antiviral effects at either entry or exit step could be manifested in our results.

It is important to note that actin may functions at multiple stages of Orsay infection. Actin forms terminal web as well as other types of cytoskeleton networks that are under constant remodeling including polymerization, depolymerization, branching, etc. We have previously shown that Orsay infection resulted in the disruption/reorganization of the terminal web in C. elegans intestinal cells for non-lytic exit of the virus (6). Likewise, an earlier study of the intracellular parasite *N. parisii* showed that gaps were formed in the terminal web of C. elegans intestinal cells when *N. parisii* meronts differentiate into spores, and the formation of gaps may remove a barrier to enable spores to exit into the lumen (32). While actin may be used by the host for antiviral defense, it can also facilitate viral infection in various ways, e.g. intracellular transport (33), long-distance dissemination of viruses (34), etc. The complex role of actin is further demonstrated by the observation that different actin remodelers can have opposing effects on Orsay infection. While we discovered here that the *C. elegans* WASP ortholog *WSP-1* and the WASP regulator *CDC-42* are required for the host antiviral defense (Fig. 4), Jiang et al reported that another *C. elegans* WASP ortholog *VIRO-2* and the WASP regulator *NCK-1* are required for virus infection (35). Consistent with this report, we also found that *nck-1(RNAi)* reduced viral load (Fig. 4F). RNAi of another WASP regulator *wip-1* also reduced viral load (Fig. 4F), suggesting that *WIP-1* shares a similar function with *NCK-1* in promoting viral infection. Our results demonstrate the complexity of host-virus interactions and provide a framework to unravel these interactions.

## MATERIALS AND METHODS

### Strains

*C. elegans* strains were maintained on standard nematode growth medium (NGM) seeded with *E. coli* OP50 as described (36). All strains were maintained at 20°C except those with *glp-4(bn2ts)* mutation, which were maintained at 15°C.

The following strains were obtained from the *Caenorhabditis* Genetics Center (CGC): N2, BE3 *sqt-2(sc3) II*, CB566 *unc-34(e566) V*, MT12615 *mys-1(n3681) V*, MT13649 *nurf-1(n4295) II*, MT15795 *isw-1(n3294) III*, NG324 *wsp-1(gm324) IV*, RB1963 *col-61(ok2582) I*, VC2706 *wve-1(ok3308) I/hT2[bli-4(e937) let-?(q782) qIs48] (I;III)*, VC898 *cdc-42(gk388)/mIn1[mIs14 dpy-10(e128)] II*, VC971 *+/mT1 II; act-5(ok1397)/mT1[dpy-10(e128)] II*.

SS104 *glp-4(bn2ts) I* (37) was kindly provided by Dr. Natasha Kirienko. FX05448 *col-51(tm5448) I* was kindly provided by Dr. Shohei Mitani. ERT54 *jyls8[Ppals-5::GFP; Pmyo-2::mCherry]* (21) was kindly provided by Dr. Emily Troemel. MW265 *rde-1(ne219);Is[Pges-1::rde-1::unc54 3’UTR,Pmyo-2::RFP]* was kindly provided by Dr. Meng Wang.

We generated the following strains: WWZ231 *col-51(tm5448) I* by outcrossing FX05448 four times, WWZ381 *col-61(ok2582) I* by outcrossing RB1963 six times, WWZ217 *drh-1(ok3495)IV;glp-4(bn2ts) I (6)*, WWZ369 *col-51(tm5448) I*; *jyls8[Ppals-5::GFP; myo-2::mCherry]*, WWZ362 *cazEx26[Pcol-51::col-51:::GFP, Pmyo-3::mCherry]* by cloning 1397bp promoter and 1266bp coding sequence of *col-51* in frame with GFP into the vector pPD95.77 (Fire Lab *C. elegans* vector kit; Addgene) and microinjecting to N2 worms.

### Viral filtration

Small amount of viral filtration was prepared as previously described (23). To scale up, two 6 cm NGM agar plates of infected *rde-1(ne219)* worms were cultured by adding 40 μl of the viral filtration to each plate. These worms, together with 2 ml of viral filtration, were then used to seed 50 ml liquid culture following the standard liquid culture protocol (36). Worms were cultured in a 20°C incubating shaker for 5 days. The culture was then cleared by centrifugation at 20,000g for 10 minutes at 4°C. The supernatant was passed through a 0.22 μm filter, and aliquots were kept at -80°C.

### RNAi screen

RNAi by feeding was performed using a method modified from standard protocol (38). Bacteria from published RNAi library (39) were cultured in 2.4 ml of L-broth containing 50 μg/ml carbenicillin on 24-well deep-well plates on a 37°C incubator shaker overnight. 1 mM Isopropyl β-d-1-thiogalactopyranoside (IPTG) was then added to the culture to induce bacteria. After two hours of induction, bacteria were pelleted by centrifugation, and culture media were discarded. The cells were then resuspended in 60 μl S medium (36) containing 50 μg/ml carbenicillin and 1 mM IPTG.

Synchronized L1 larvae from the strain SS104 were obtained by bleaching (36). ∼100 L1s were placed in each well of a 96-well plate with 30 μl of RNAi bacteria suspension, and the volume was brought up to 100 μl with S medium with 50 μg/ml carbenicillin and 1 mM IPTG. For the virus-infected group, 1 μl of Orsay viral filtration was added to each well. The plates were parafilmed and placed on a 20°C incubator shaker for five days till the worms reached day-3 adults. Worms were then transferred to unseeded scanning plates (modified NGM plates that do not contain peptone or cholesterol), killed by sodium azide, and observed under a high-contrast stereoscope for the transparent intestine phenotype.

A quick manual inspection was applied to select RNAi wells that showed qualitative differences between infected and uninfected groups. The worms on these wells were then quantitatively scored. The RNAi effects of hit genes were then confirmed in at least three independent experiments. The identity of hit genes in the RNAi bacterial clones was confirmed by Sanger sequencing.

### Small-scale experiments for scoring infection symptom of transparent intestine

Standard 6-cm NGM agar plates seeded with OP50 were prepared as described (36). For RNAi experiments, RNAi plates (38) were used. To make RNAi plates, RNAi bacteria were cultured in L-broth containing 50 μg/ml carbenicillin at 37°C overnight. Bacteria were then used to seed 6-cm NGM agar plates that contained 50 μg/ml carbenicillin and 1 mM IPTG. *act-5* RNAi bacteria was diluted 1:25 with control empty vector bacteria before seeding. The seeded RNAi plates were left at room temperature overnight.

∼100 synchronized L1 larvae were dropped onto each plate. For the infected group, 40 μl of viral filtration was added to each plate. The worms were cultured in a 20°C incubator for five days until they were day-3 adults and scored. For strains that did not have the *glp-4(bn2ts)* mutation, adult worms were manually picked and transferred to a fresh RNAi plate everyday once they reached the day-1 adult stage to avoid interference from progeny.

### RNA quantification by qRT-PCR

For standard culture, ∼500 synchronized L1 larvae were dropped onto each 6-cm NGM agar plates seeded with OP50 (36). 40 μl of viral filtration was added to each plate in the infected group.

For RNAi worms, 10 L4 animals were placed on each well of a 6-well RNAi plate seeded with RNAi bacteria, and cultured in a 20°C incubator for 24 hours. The worms were then removed to keep only synchronized eggs on the plates. 20 μl of viral filtration was added to each well for the infected group.

The worms were cultured in a 20°C incubator for two days till they reached L4 stage. L4 worms were collected and washed four times with 10 ml S basal medium (36). Total RNA from worms was extracted by using TRIzol (Invitrogen), digested with DNase (Invitrogen), and reverse transcribed to cDNA using RETROscript (Thermo). qRT-PCR was performed by using PerfeCTa SYBR green SuperMix (Quantabio). Primer pairs GW194/GW195 and AMA-1F/AMA-1R (23) were used to target Orsay virus RNA1 and the internal reference gene *ama-1*, respectively. Primer pair WSP-1 QF (ACGACGATGATATGGATGAAGC) and WSP-1 QR (GTATTGACGCACTGGTGTTTGA) was used to quantify *wsp-1* RNA level. The quantification cycle (Cq) values of Orsay or *wsp-1* were normalized to that of *ama-1*. Three technical replicates were tested in each trial. At least two independent trials were performed for each test group.

### Chemical treatment

20 mM stock solutions of bismuth subsalicylate (Sigma-Aldrich 480789) and resorcinol monoacetate (Sigma-Aldrich R856) were made by dissolving chemicals in the solvent dimethyl sulfoxide (DMSO). OP50 bacteria were cultured in L-broth overnight at 37°C, then pelleted and resuspended in S medium at 2.5% of the original culture volume.

For the “all time” group, ∼100 synchronized L1 worms of the strain WWZ217 were placed in each well of a 96-well plate with 30 μl of bacteria suspension, 1 μl of viral filtration, and 0.5 μl of 20 mM chemical stock solution (or 0.5 μl DMSO control). The volume was then brought up to 100 μl with S medium. The plates were parafilmed and placed on a 20°C incubator shaker for five days.

For the “post exposure” group, L1 worms were placed in each well with bacteria, viral filtration, and S medium. The testing chemicals were not added at this time. The worms were incubated on a 20°C incubator shaker for 24 hours. Then they were washed four times with S medium to remove the virus. After washing, 30 μl of bacteria suspension and 0.5 μl of 20 mM chemical stock solution (or DMSO control) were added to each well, and the volume was brought up to 100 μl with S medium. The plates were parafilmed and placed on a 20°C incubator shaker for four more days.

When the worms reached day-3 adults, they were transferred to unseeded scanning plates, killed by sodium azide, and observed under a high-contrast stereoscope for the transparent intestine phenotype.

### Infection dynamics

About 100 synchronized L1 larvae of ERT54 or WWZ369 were placed on a 3-cm NGM agar plate seeded with OP50, and cultured at a 20°C incubator for 48 hours till they reached L4 stage. A mixture of 15 μl water and 5 μl orsay was dropped on the plate and the timer was set as time point 0. Worms were observed for GFP under a Zeiss SteReo Discovery V20 stereoscope starting from time point 14 hours to time point 40 hours. GFP-positive worms were counted and removed from plates. A control group of uninfected worms was always used as a negative control to ensure that no GFP was observed in those worms.

### Microscopy

Epifluorescent images were taken using a Zeiss AxioImager M2m microscope equipped with a Zeiss AxioCam MRm camera and AxioVision software 4.8.

### Quantitative Epistasis

Quantitative epistasis analysis was performed using math model as previously described (40). At least three independent trials were performed for each gene pair. In each trial, at least two plates were tested for each genotype.

## ACKNOWLEDGEMENTS

We thank Drs. Natasha Kirienko, Shohei Mitani, Emily Troemel, and Meng Wang for reagents; Jim Zhang for critical reading of the paper; and Joaquina Nunez for technical support. We thank the Caenorhabditis Genetics Center (CGC), which was funded by the NIH Office of Research Infrastructure Programs (grant P40 OD010440), for providing strains. This work was supported by the Robert A. Welch Foundation (C-1565 to Y.J.T.), the Hamill Foundation (the Hamill Award at Rice University to Y.J.T. and W.Z.), and the National Institutes of Health (R01-AI122356 to Y.J.T. and W.Z; R21-AI171624 to Y.J.T.). The funders had no role in study design, data collection and analysis, decision to publish, or preparation of the manuscript.

